# Community RNA-Seq: Multi-kingdom responses to living versus decaying root inputs in soil

**DOI:** 10.1101/2021.01.12.426429

**Authors:** Erin E. Nuccio, Nhu H. Nguyen, Ulisses Nunes da Rocha, Xavier Mayali, Jeremy Bougoure, Peter Weber, Eoin Brodie, Mary Firestone, Jennifer Pett-Ridge

## Abstract

Roots are the primary source of organic carbon inputs to most soils. Decomposition is a multi-trophic process involving multiple kingdoms of microbial life, but typically microbial ecology studies focus on one or two major lineages in isolation. We used Illumina shotgun RNA sequencing to conduct PCR-independent SSU rRNA community analysis (“community RNA-Seq”) to simultaneously study the bacteria, archaea, fungi, and microfauna surrounding both living and decomposing roots of the annual grass, *Avena fatua*. Plants were grown in ^13^CO_2_-labeled microcosms amended with ^15^N-root litter. We identified rhizosphere substrate preferences for ^13^C-exudates versus ^15^N-litter using NanoSIMS microarray imaging (Chip-SIP). When litter was available, rhizosphere and bulk soil had significantly more Amoebozoa, which are potentially important yet often overlooked top-down drivers of detritusphere community dynamics and nutrient cycling. Bulk soil containing litter was depleted in Actinobacteria but had significantly more Bacteroidetes and Proteobacteria. While Actinobacteria were abundant in the rhizosphere, Chip-SIP showed Actinobacteria preferentially incorporated litter relative to root exudates, indicating this group’s more prominent role in detritus elemental cycling in the rhizosphere. Our results emphasize that decomposition is a multi-trophic process involving cross-kingdom interactions, and the trajectory of carbon through this soil food web likely impacts the fate of carbon in soil.

## INTRODUCTION

Soil carbon is derived primarily from decomposed plant material (1, 2) and the fluxes that control the size of this pool are critical to the global carbon (C) cycle. The soil adjacent to plant roots (the rhizosphere) is a nexus for root C input, microbial C transformation, as well as C loss through decomposition (3, 4). Most root C is decomposed to CO_2_, and the remainder typically undergoes multiple microbial transformations before it is stabilized. The spatial organization of soil habitats such as the rhizosphere and detritusphere (a region containing non-living organic matter) is particularly important for carbon and nutrient transfer by soil microbes and fauna, and the characteristics and rates of these microbial transformations determine how much carbon remains in soil (5). While it is widely recognized that fungi and soil fauna are instrumental to decomposition (6), less is known about how the greater soil food web of bacteria, archaea, fungi, and microfauna responds to decomposing litter in the rhizosphere and detritusphere.

To date, microbial ecology surveys studying litter decomposition using amplicon sequencing have primarily focused on bacteria or fungi, but decomposition is conducted by a broad array of organisms (7) including microfauna (here we use this umbrella term to include protists, nematodes and other soil invertebrates < 100 μm) (6, 8). Fungi play a key role in the decomposition of plant litter by providing the majority of the extracellular enzymes needed to depolymerize plant residues (9) and have been well-studied (10–13). Litter-associated microfauna may consume and directly break down litter (6), and protists and nematodes are also known to consume fungi and bacteria (8, 14-17); the presence of these consumers can affect both the microbial community composition and the rate of litter decomposition (18–24). Improving our understanding of decomposition in soil necessarily requires us to consider the roles and trophic interactions among the broader soil food web.

In the past decade, amplicon metabarcoding with high-throughput sequencing approaches have allowed the identification of multiple groups of soil organisms in parallel (25–27). However, PCR amplification has multiple layers of biases, including primer selection and bioinformatic processing, and the lack of a universal primer set means multiple primer sets are required to amplify taxonomically disparate groups (28–30). An alternative approach is to use amplification-independent methods, such as shotgun RNA sequencing (RNA-Seq) for community analysis, which we call “community RNA-Seq”. This method not only reduces the inherent biases associated with PCR (31–33), but since rRNA is an integral part of ribosomes that controls protein synthesis across multiple kingdoms of life (34), direct sequencing of RNA allows us to study active communities within Bacteria, Archaea, and Eukarya simultaneously without amplification. Additionally, as most RNA is ribosomal RNA, the resulting sequences have naturally high coverage of the ribosomal subunits most frequently used for taxonomic analysis (e.g., 16S, 18S, 28S) (35), which allows greater sequencing depth of taxonomic markers than metagenomic sequencing. To assess community composition, community RNA-Seq is followed by either reassembling rRNA fragments into full ribosomal subunits or classifying the short reads directly (35–38). Community RNA-Seq has been used to a limited degree in microbial ecology due to the difficulty working with RNA in an environment such as soil, but initial studies suggest it is a particularly useful approach to study protists in soil without PCR and cultivation biases (35, 39). Eukaryotic primers are not universal for protists (40), which has led to Amoebozoa being underrepresented in SSU rRNA gene surveys due to long SSU regions, and ciliates being overrepresented due to short SSU regions (39).

Methods that leverage isotopes as tracers of microbial activity (e.g. assimilation of substrates) are capable of adding another layer of ecological information to community analysis studies and have the potential to expand our understanding of food web dynamics and nutrient cycling in multi-trophic communities. Stable isotope probing (SIP) is a suite of powerful methods to study microbial ecophysiology in complex environments (41, 42) where a normally rare stable isotope (e.g., ^13^C, ^15^N, ^18^O) is added to the environmental matrix, and organisms that incorporate the labeled substrate become isotopically enriched over time in proportion to their activity (43, 44). Nucleic acid-SIP techniques (43, 45) are currently the most widely used means to directly connect microbial identity to substrate utilization. An iteration of nucleic acid-SIP is Chip-SIP where an imaging mass spectrometer (NanoSIMS) is used to determine the isotopic enrichment of rRNA hybridized to a phylogenetic microarray (46, 47). This method has a low ^13^C enrichment requirement (0.5 atom%) relative to standard SIP, permits shorter isotope incubations, allows dual ^13^C and ^15^N labels in the same sample, and requires no amplification. Using this method, we can trace the fate of ^13^CO_2_ after it is fixed by plants and released as ^13^C-rhizodeposits, and simultaneously trace detritus labeled with another isotope (e.g., ^15^N) to examine the relative incorporation of rhizosphere versus detritus in the same community.

In this work, we used shotgun RNA sequencing and Chip-SIP to study how living versus detrital root material alters the bacterial, archaeal, fungal, and microfaunal communities in the *Avena fatua* rhizosphere and surrounding bulk soil. We hypothesized that 1) the detritusphere would increase saprotrophic fungi and eukaryotic grazers in the rhizosphere, and 2) rhizosphere decomposers would consume both litter and root exudates, rather than specialize on either resource alone.

## METHODS

### Microcosm setup and soil collection

Soils were collected at the Hopland Research and Extension Center (HREC, GPS 38.992982, - 123.067562) in Hopland, CA (USA), which experiences a Mediterranean climate (48). Soils are a fine loam Alfisol complex (Ultic Haploxeralf mixed with a Mollic Palexeralf) with 1.7% C and 0.14% N (49). The top 10 cm of soil was collected from beneath a stand of naturalized *Avena barbata* within a wild annual grassland community at 1 m intervals along a 10 m transect in January. Large plant material was removed, including root pieces, and soil was sieved to 2 mm, homogenized, then mixed with sand (1:1 w/w sand:dry weight soil) to improve drainage. The mixed soil was packed into the main chamber of plastic microcosms (15 cm × 5 cm × 40 cm) to a density of 1.2 g/cm^3^ as previously described (Fig. 1a) (50, 51). *Avena fatua* seeds (Pacific Coast Seed Inc., Tracy, CA, USA) were germinated in the dark for 7 days. One seedling per microcosm was planted once the roots were greater than 1 cm long and after the shoot had emerged from the seed. Plants were grown in a greenhouse under a 14-hour photoperiod and watered every 2-3 days to field water-holding capacity (approximately 50% saturation). After 6 weeks, the solid divider separating the main chamber from the sidecar was replaced with a slotted divider (slots ca. 10 cm x 4 mm) and the sidecar (5 mm deep) was filled with the experimental soil (Fig. 1a).

**Figure 1:**
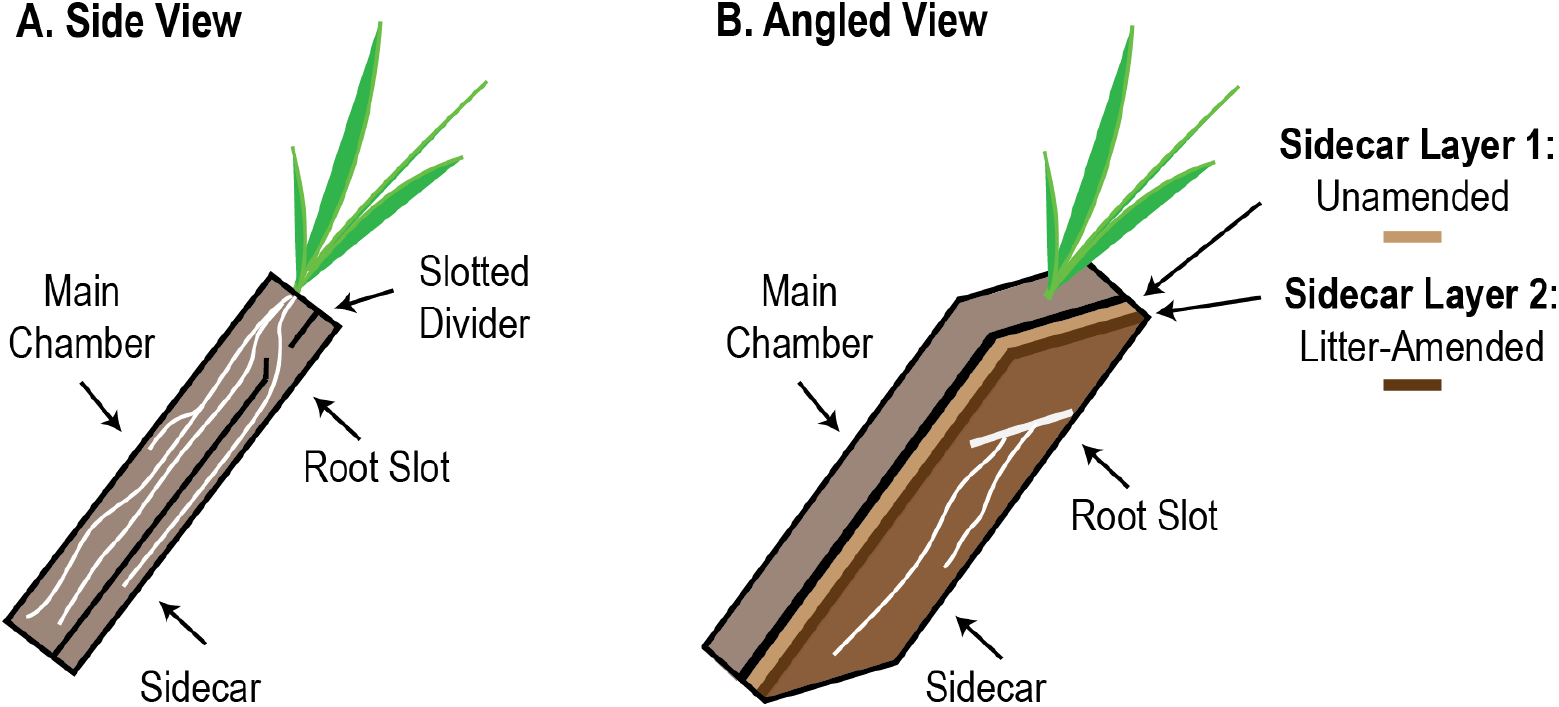
Microcosm design and sampling strategy. (A) Microcosms had a main chamber that housed the plant during plant growth and maturation. The main chamber was separated from an auxiliary root chamber (the sidecar) by a solid divider; microcosms were tilted to promote the growth of roots. After 6 weeks, the solid divider was removed and replaced with a slotted divider to permit root growth into the sidecar, and the sidecar was then filled with the experimental soil. (B) Half of the experimental soils were amended with ^15^N-labeled root detritus (layer 2), which was placed on top of unamended soil (layer 1). Unamended controls were prepared in the same manner, but no litter was added to the second layer. After 6, days the roots entered the sidecar, and the plants were then pulse labeled for 3 days with ^13^CO_2_ and harvested. Roots grew along the face of the sidecar and rhizosphere soil (<2 mm from root) and bulk soil (>4 mm from root) were excised with a scalpel.

Sidecar experimental soil was freshly collected and sieved HREC soil (not mixed with sand), and half of the microcosms received ^15^N-labeled *A. fatua* root litter chopped to ca. 1 mm (78 atom % ^15^N; see Supplemental Methods for details regarding production of this material). The ^15^N isotopic tracer allowed us to use mass spectroscopy to detect the communities that were actively consuming litter-N. For the litter treatment (n=3), the soil was added to the sidecar in two layers, each approximately 2.5 mm deep: the bottom layer contained 75 g of soil with no litter, while the top layer contained the 75 g of soil amended with 0.4 g of ^15^N root litter. For the no-litter control (n=3), 150 g of soil was added to the sidecar. After packing the sidecars, microcosms were tilted at 45° to encourage root growth into the sidecars.

After filling the sidecar, plants were grown for 6 days in prior to ^13^CO_2_ labeling, which is the amount of time it typically takes for roots to enter the sidecar. A 1.5m × 1.5m × 0.76 m plexiglass glovebox (Coy Laboratory Products, USA) was used as a labeling chamber at the UC Berkeley EPIC facility (49). The maximum chamber temperature was cycled between 26-28 °C during the day and allowed to cool naturally to 20-22 °C at night. Before dawn each day, the air in the chamber was cycled through a desiccator filled with soda lime to remove CO_2_ until the chamber atmosphere reached < 25 ppm CO_2_. The chamber was then filled with 99 atom % ^13^CO_2_ until the concentration reached a set point of 400 ppm, and maintained at 400 ppm throughout the day using an SBA-5 model IRGA (PP Systems, 400 ppm ^13^CO_2_ standard calibration) attached to a CR800 model datalogger (Campbell Scientific, Logan, UT, USA). Using this setup, the plants were labeled with ^13^CO_2_ for 3 days.

After the 3 days of labeling, the front plates of the sidecars were removed to access an intact rhizosphere along the entire length of a root. Rhizosphere soil within 2 mm of the roots was excised using a scalpel. The soils were immediately placed in ice-cold Lifeguard RNA protectant solution (MoBio). Tubes were shaken for 2 minutes on a horizontal vortex adaptor (MoBio) on medium speed to release soil from the roots. The tubes were centrifuged at 2.5 × *g* for 1 min at 4°C, and any roots or floating root litter were removed with flame-sterilized forceps. The remaining soil was pelleted by centrifuging at 2.5 × *g* for 5 min at 4°C. After the supernatant was carefully removed, the pellets were immediately frozen on dry ice, and stored at −80°C for molecular analysis. Soil >4 mm from a root was treated as bulk soil. To collect bulk soils with litter, the top half of the sidecars that contained ^15^N labeled litter was randomly excised using a scalpel. These samples often contained visible pieces of ^15^N labeled roots. Bulk soil samples were processed in the same manner as rhizosphere soils. We collected a total of 12 soil samples: 2 locations (rhizosphere, bulk) × 2 litter conditions (litter, no litter) × 3 replicate microcosms. Hereafter, we refer to samples from the unamended control as “rhizosphere” and “bulk” and samples from the litter addition treatment as “rhizosphere-litter” and “bulk-litter”.

### RNA Extraction and Sequencing

RNA was extracted in triplicate from 0.2 g soil per sample using the phenol-chloroform extraction protocol (52), modified from (53). Extracted nucleic acids were passed through the Allprep DNA/RNA Mini Kit (Qiagen Sciences, Maryland, USA) to separate RNA from DNA. RNA was treated with DNase using an on-column DNase digestion. For community RNA-Seq, metatranscriptomic libraries were prepared directly from total RNA without rRNA removal using the TruSeq RNA Kit (Illumina, Inc., San Diego, CA, USA) according to the manufacturer’s instructions. Metratranscriptomic libraries were sequenced on an Illumina GAIIX sequencer using 150 basepair (bp) paired-end sequencing at Lawrence Berkeley National Laboratory with an average of 9.5 million paired reads per sample.

### Sequence quality control and rRNA reconstruction

Sequences were demultiplexed, and sequence quality was checked with FastQC (54). We used Trimmomatic (55) with default parameters with one exception; we removed the first 10 bp from the 5’ end due to overrepresentation of this region in the dataset. Sequences shorter than 60 bp after trimming were removed. Reads that did not pair were discarded.

EMIRGE (36) was used to reconstruct near-full-length SSU rRNA sequences for Bacteria, Archaea, and Eukarya using the script “emirge_amplicon.py”. The script was run on paired-end reads with the following parameters: mean insert length of 342, insert standard deviation of 100, and max read length of 151. The Greengenes 13_5 database clustered at 97% similarity was used to create the reference database for Bacteria and Archaea (56). The SILVA 114 NR database (57) was used to create the reference database for Eukarya. The database was also clustered at 97% as above. After the databases were created, the non-standard characters were altered as previously described (36). Bowtie indices required by EMIRGE were calculated for the databases using bowtie-build (58).

### OTU clustering and classification

Bacterial and archaeal sequences were analyzed separately from eukaryotic sequences. Sequences were clustered using UPARSE (usearch_v7) (59) and analyzed using QIIME 1.8 (60) at 97% sequence similarity. OTUs were classified using the RDP classifier (61), where bacterial and archaeal classifications were trained using Greengenes 13_5 and eukaryotic sequences were trained using SILVA 119NR (57). UCHIME (62) was selected to detect chimeras after testing three chimera-checking tools (see Supplemental Methods). OTUs were required to be present in at least two samples, and OTUs classified as chimeras or plant and algal chloroplasts were removed from the dataset. In total, we analyzed 7229 unique full-length bacterial and archaeal RNA sequences created by EMIRGE (1127 OTUs at the 97% similarity level), and 8488 unique full-length eukaryotic RNA sequences created by EMIRGE (265 OTUs at 97% similarity level).

Since EMIRGE calculates a relative abundance estimate for each consensus sequence, a custom OTU table (sample × OTU matrix) was created after OTU picking to incorporate relative abundances of the consensus sequences into the microbial community analysis. To convert the consensus sequence relative abundance into sequence abundance, we multiplied the total number of reads that Bowtie mapped to the database by the relative abundance derived from the “normalized priors”, as per Miller, Handley (63): total mapped reads × consensus sequence relative abundance = number of sequences per consensus sequence. Since each OTU can contain multiple consensus sequences, we calculated the OTU sequence abundance by summing the number of sequences for each consensus sequence within the OTU. The samples were then rarefied to an even depth of 121 737 sequences for bacteria and archaea, and 27 668 sequences for eukaryotes. As per the recommendations of Miller, Handley (63), OTUs with less than 0.01% relative abundance were removed.

### Statistical Analysis

Community differences were visualized by Principal Components Analysis (PCA) in QIIME using a pairwise weighted Unifrac distance matrix (64). To determine which OTUs differed in relative abundance between the litter and unamended treatments, we performed two sets of parametric t-tests in QIIME (group_significance.py): rhizosphere vs. rhizosphere-litter; bulk vs. bulk-litter. Only OTUs that were detected in all three replicates of at least one treatment were considered for analysis. P-values were corrected for multiple comparisons using an FDR correction. To calculate kingdom-or phylum-level relative abundances, relative abundances were summed for all OTUs within each group (kingdom for Eukarya, phyla for Bacteria and Archaea) and significant differences were determined using a t-test. Changes in the relative abundances for each group were determined by comparing litter-amended samples to their unamended control for bulk and rhizosphere soil separately (i.e., bulk vs. bulk-litter; rhizosphere vs. rhizosphere-litter).

### Chip-SIP Analysis

To follow the movement of C and N from living plants and dead roots into the microbial community, we analyzed the rhizosphere of a microcosm containing both ^15^N litter and ^13^C exudates using Chip-SIP, a method that can detect and quantify ^15^N/^14^N and ^13^C/^12^C ratios of labeled RNA hybridized to microarrays (46, 47). Detailed methods for probe design, microarray synthesis and hybridization, NanoSIMS analysis, and data processing can be found in the Supplemental Methods. Briefly, we designed a microarray with probes using ARB (65) for the 180 most abundant Bacteria, Archaea, and Eukarya (fungi, protists, nematodes) OTUs found in this study, as well as probes targeting plant chloroplasts; a taxonomy summary of the probes used in this study is available in Table S1. Ten distinct probes per OTU were printed in three replicate blocks on the microarray. We produced 2 microarrays for this sample, one to detect RNA binding, and a second to detect RNA ^13^C and ^15^N isotopic enrichment with NanoSIMS. To detect RNA binding, RNA was labeled with Alexafluor 532 dye using the Ulysis kit (Invitrogen), fragmented with fragmentation buffer (Affymetrix), purified, concentrated and hybridized onto the first array. For NanoSIMS analysis, unlabeled RNA was again fragmented, purified and concentrated and then hybridized to the second array. The array with fluorescently labeled RNA was imaged with a Genepix 4000B fluorescence scanner. The second array (with non-fluorescently labeled RNA) was also imaged with the fluorescence scanner to allow navigation to analysis spots in the NanoSIMS. Data were collected on the LLNL NanoSIMS 50 in pulse counting mode using aperture slit 3 and entrance slit 5, first collecting ^12^C^14^N^-^ and ^12^C^15^N^-^, and then ^12^C^14^N^-^ and ^13^C^14^N^-^. The resulting data were visualized as a stitched isotope map (Fig. S1) and data extracted as per Mayali, Weber (46).

The proportion of isotopes is presented as a relative atom percent excess (APE) enrichment ratio of ^13^C to ^15^N (^13^C-APE:^15^N-APE) to indicate substrate preferences, where lower values indicate greater ^15^N enrichment in the RNA, and higher values indicate greater ^13^C enrichment in the RNA. Due to the higher background of ^13^C compared to ^15^N on the array, we used a normalization factor of 1.7 to calculate these relative enrichment ratios, as previously described (66). Higher relative enrichment in ^15^N is interpreted as having a preference for amended ^15^N root litter whereas higher relative enrichment in ^13^C is interpreted as having a preference for ^13^C root exudates. We note that this is a relative measure as the ^13^C values do not reflect the total ^13^C ingested, as part of the ^13^C consumed is lost through respiration (66).

## RESULTS

### Community Structure from Reconstructed SSU rRNA

The addition of root litter and the presence of living roots both significantly altered the bacterial and eukaryotic communities relative to bulk soil; both the bacterial and eukaryotic communities had significantly different clusters per treatment by PERMANOVA analysis (Fig. 2) (see Table S2 for F Tables), though the eukaryotic communities had more overlap (Fig. 2b). The bulk-litter communities were the most distinct group in ordination space for both bacteria and eukaryotes. Root litter had the strongest effect on both the bacterial and eukaryotic communities, explaining 30% and 28% of the variance in those communities, respectively (2-way PERMANOVA: bacteria F_1,4_ = 7.2, r^2^ = 0.30, p > 0.001; eukaryotes F_1,4_ = 5.4, r^2^ = 0.28, p > 0.001). The presence of a living root also significantly altered these communities, which was strongly significant for bacteria (2-way PERMANOVA: F_1,4_= 4.7, r^2^ = 0.20, p = 0.006), and was slight but significant for eukaryotes (2-way PERMANOVA: F_1,4_ = 3.2, r^2^ = 0.17, p = 0.029).

**Figure 2:**
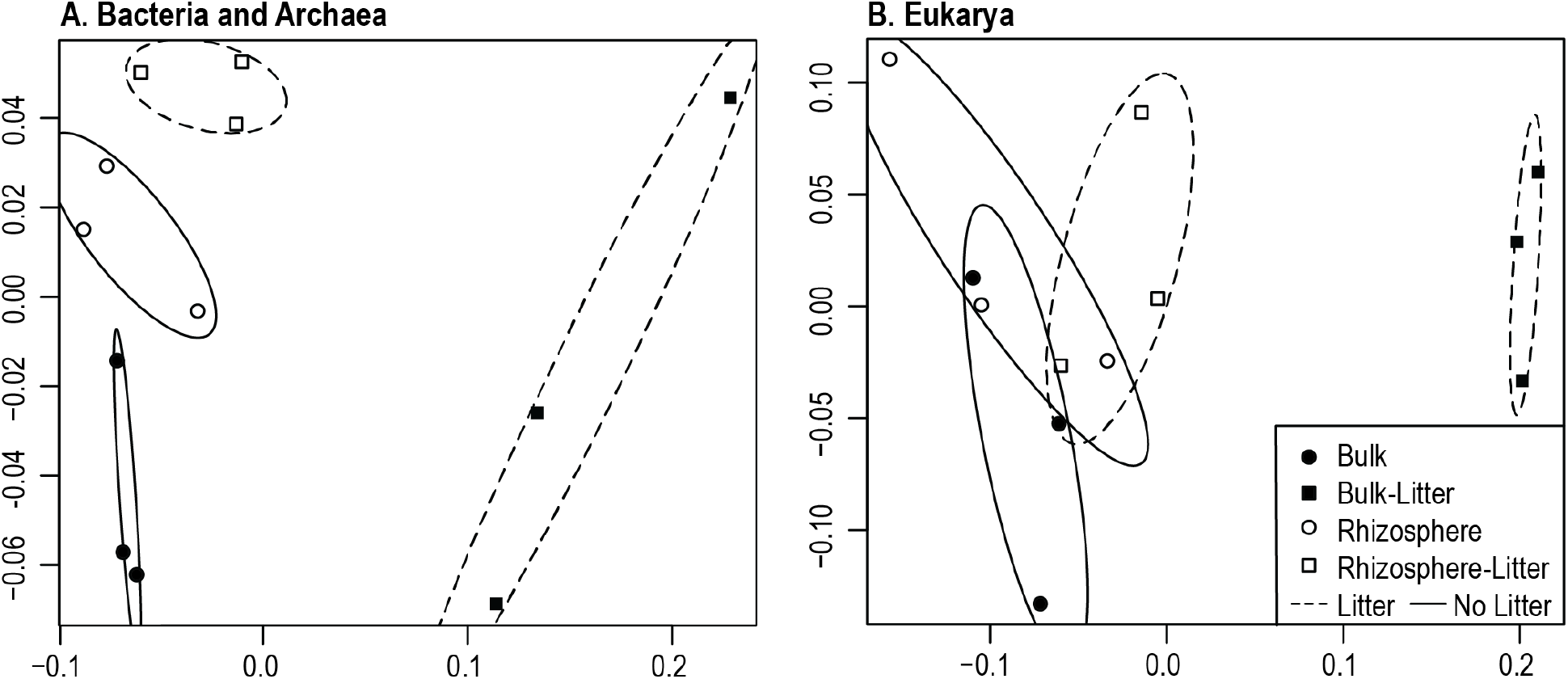
Community RNA-Seq non-metric multidimensional scaling ordinations for (A) Bacteria and Archaea (assembled 16S rRNA), and (B) Eukarya (assembled 18S rRNA) in the *Avena fatua* rhizosphere and bulk soil in response to litter amendments. Soil was sampled after 3 days of fresh root growth into the microcosm sidecars. Filled symbols represent bulk soil, while hollow symbols represent rhizosphere soil. Squares indicate litter addition while circles had no litter added. Ovals represent the 95% standard error of the weighted average of scores per group (r package: ordiellipse) for litter treatments (dashed lines) and no litter (solid lines).

### Phylum and Kingdom Level Responses

We observed broad patterns in relative abundance of bacteria and eukaryotes at the phylum and kingdom level, respectively. Proteobacteria and Actinobacteria were the most abundant bacterial groups in the rhizosphere (Fig. 3a). Actinobacteria, Acidobacteria, and Chloroflexi were significantly reduced in the bulk-litter treatment (t-test: p < 0.05) (Fig. 3a), while Bacteroides and Proteobacteria were significantly more abundant in the bulk-litter treatment. For the eukaryotes, Amoebozoa were significantly more abundant in presence of litter in both rhizosphere and bulk soils compared to their respective unamended controls (Fig. 3b). In unamended bulk soil, Rhizaria were significantly more abundant. While the litter-containing treatments appear to have less Fungi, these differences were not significant (p > 0.5) compared to the unamended controls.

**Figure 3:**
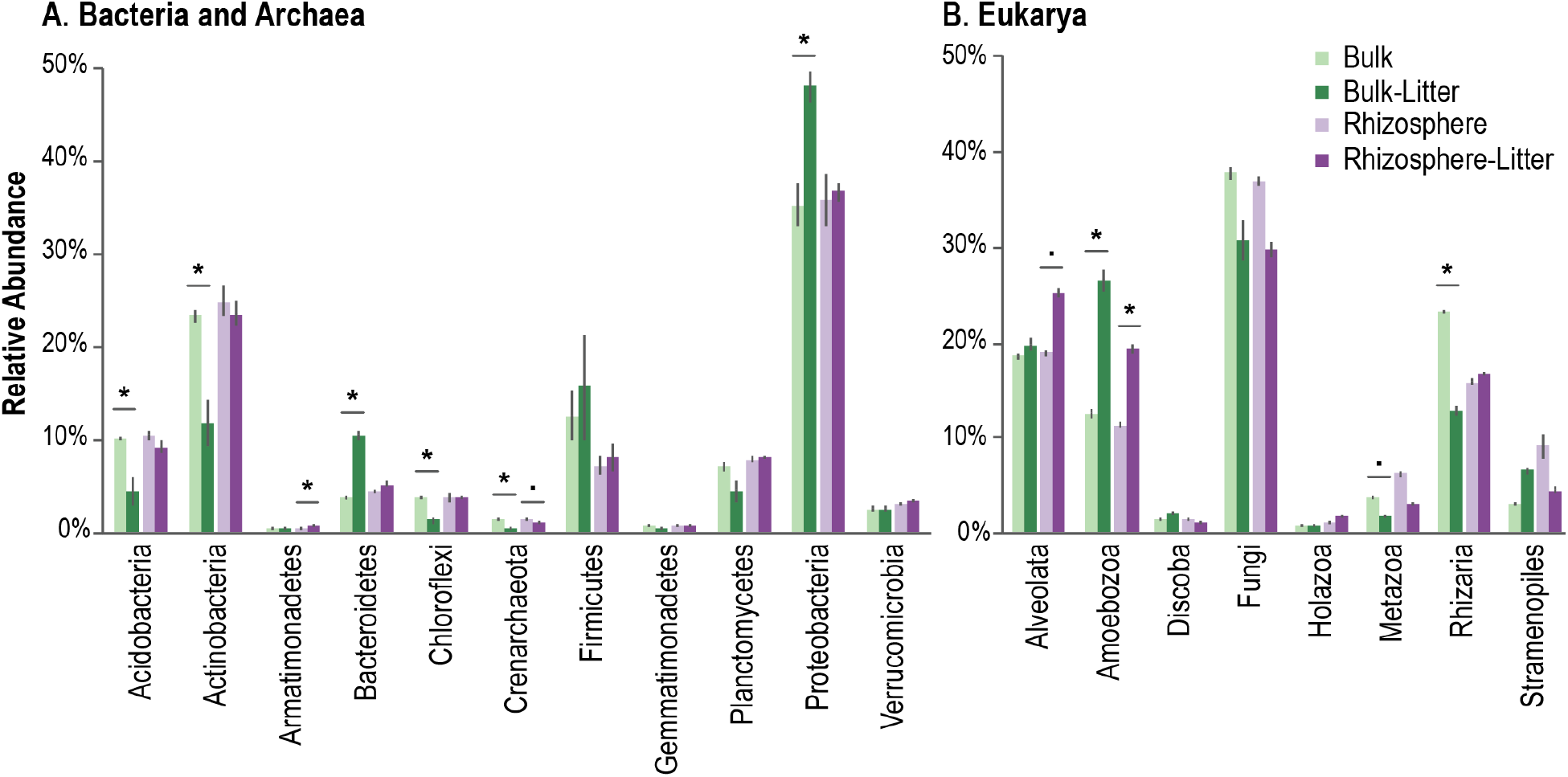
SSU rRNA relative abundance at the (A) phyla level for Bacteria and Archaea, and (B) kingdom for Eukarya. The treatments were as follows: bulk soil with no litter amendment (bulk, light green) and amended with root litter (bulk-litter, dark green), and rhizosphere soil with no litter amendment (rhizosphere, light purple) and amended with root litter (rhizosphere-litter, dark purple). Groups that significantly differed in relative abundance with litter amendments are indicated by * (t-test: p < 0.05) (Bulk vs. Bulk-Litter; Rhizosphere vs. Rhizosphere-Litter), and “.” indicates marginal significance (p < 0.1).

### Significant Litter and Rhizosphere Responders

We compared litter-amended soil to the unamended control for bulk soil and the rhizosphere (Fig. 4). In bulk soil, litter additions significantly increased specific groups of protists, fungi, and bacteria, whereas litter amendments in the rhizosphere had fewer significant responders. Protists from multiple lineages were more abundant in the presence of litter (Fig. 4b), where *Colpoda* sp. (Alveolata), *Glaseria* sp. (Amoebozoa), and *Naegleria* sp. (Heterolobosea) were some of the most abundant genera (Fig. 4b). Platyophyra sp. (Alveolata) were abundant in the rhizosphere and bulk soil when litter was present. Fungal saprotrophic *Chaetomium* sp. (Ascomycota) responded the most strongly to litter, while the other fungal taxa were more abundant in the absence of litter (*Cryptococcus* sp., *Davidiella* sp.). The bacterial taxa that strongly responded to the litter were *Massilia* sp. in the Oxalobacteriaceae (Proteobacteria), and OTUs in the families Paenibacilliaceae (Firmicutes), and Sphingobacteriaceae (Bacteroidetes) (Fig. 4a). When the rhizosphere was amended with litter, only bacteria in the family Sphingobacteriaceae (Bacteroidetes) significantly increased. Detailed taxonomic results can be found in Table S3.

**Figure 4:**
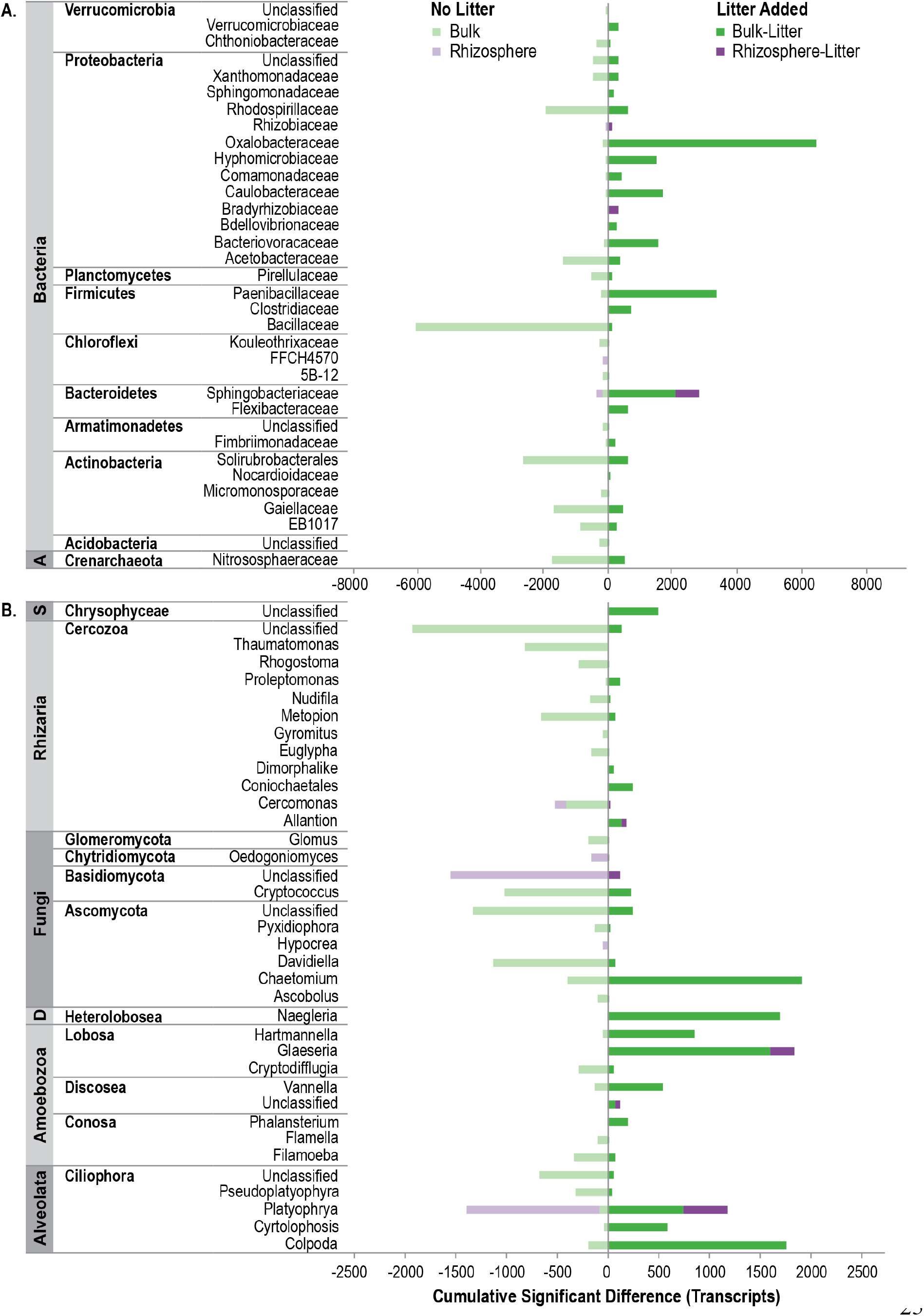
Cumulative significantly-different transcripts for positive and negative responders to detrital root litter in bulk soil and the rhizosphere. The effects of litter amendments were calculated separately for bulk soil and rhizosphere soil, where “dark green” indicates cumulative positive responses to litter for bulk soil, and “dark purple” indicates cumulative positive responses to litter for rhizosphere soil. Negative responses to litter (or preference for unamended soil) are “light green” for bulk soil and “light purple” for rhizosphere soil. Transcripts were aggregated by taxonomic family for Bacteria and Archaea and genus for Eukarya; OTU transcript abundances were averaged across replicates (n=3) prior to aggregation. Multiple comparisons were accounted for using a FDR p-value correction.

When no litter was present, an unclassified fungus in the phylum Basidiomycota and *Platyophrya* sp. (Alveolata) responded strongly to the rhizosphere. Protists from the Rhizaria, (phylum Cercozoa) were more abundant in unamended soil, particularly unclassified genera within the classes Thicofilosea and Eugliphida. Taxa that were more prominent in unamended bulk soil were from the Rhodospirillaceae (Proteobacteria), Bacillaceae (Firmicutes), Solirubrobacterales (Actinobacteria).

### Chip-SIP: Substrate Preferences

We used Chip-SIP stable isotope analysis to distinguish substrate preferences in bacterial, archaeal, and eukaryal taxa between root exudates (^13^C enriched) and decaying root litter (^15^N enriched). In the ^13^C-labeled rhizosphere sample amended with ^15^N-litter, we detected 42 isotopically enriched OTUs with Chip-SIP (Fig. 5; 1 archaea, 33 bacteria, 8 fungi); protists probes were on the array but were not enriched during this 3-day ^13^CO_2_ experiment (see probe taxonomy in Table S1). No microorganism’s RNA was enriched solely in ^13^C or ^15^N, and only the plant probes on the array were solely enriched with ^13^C and had the highest relative enrichment ratios of the dataset (Table S4). As a phylum, the Actinobacteria OTUs contained a relatively higher proportion of ^15^N than ^13^C: 6 of 7 enriched taxa fell on the lower range of the ^13^C/^15^N spectrum (0.2 – 0.8). Xanthobacteriaceae bacteria, *Sphingomonas* bacteria, and Eurotiomycetes fungi were also more enriched in ^15^N relative to the other Chip-SIP taxa (Table S4), however RNA of *Sphingomonas* sp. and Eurotiomycetes were also equally or more enriched in ^13^C, suggesting that these organisms consumed both fresh and detrital plant material during this 3-day study. Organisms with preference for rhizosphere exudates were 2 *Bacillus* OTUs, 2 Rhizobiales OTUs (Bradyrhizobiaceae, *Rhizobium*), and 2 Burkholderiales OTUs (*Massilia*). Fungi from the Ascomycota (Dothideomycetes, Eurotiomycetes, Leotiomycetes) and Basidiomycota (Agaricomycetes) tended to be equally or slightly more relatively enriched in ^13^C than ^15^N.

**Figure 5.**
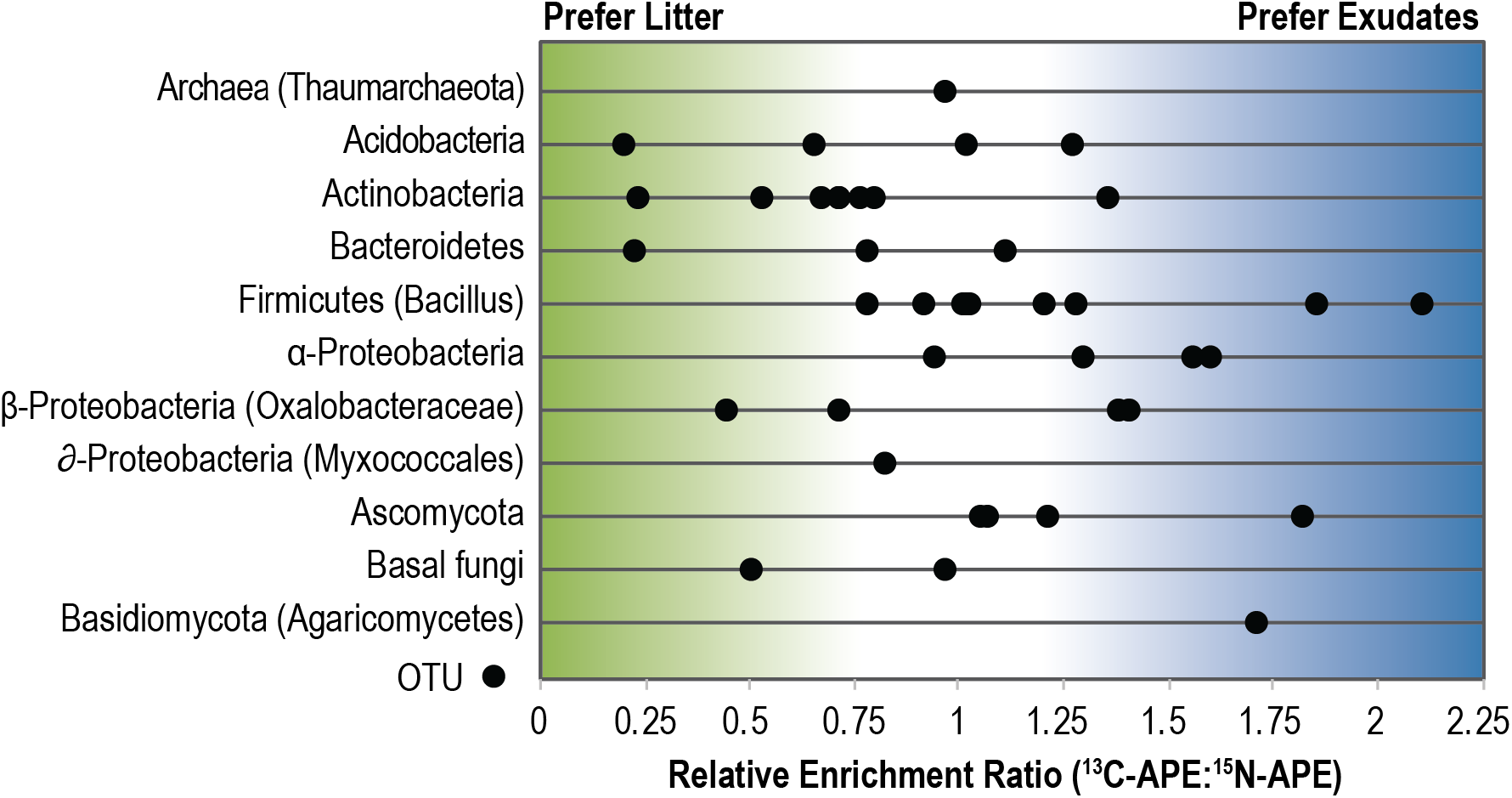
Relative substrate preferences for Bacteria, Archaea, and Fungi detected by Chip-SIP. Each dot represents a taxon that was significantly enriched in ^13^C or ^15^N derived from ^13^C-exudates or ^15^N-root detritus. The x-axis is the ratio of the atom percent excess (APE) ^13^C enrichment and ^15^N enrichment for the probe set, which is a unitless relative measure; the ratio was corrected by 1.7 to account for dilution of the C signal by the chip surface. The position of the taxon along the x-axis indicates its preference for exudates or root litter. Those that are positioned towards the left (green) incorporated relatively more isotope from ^15^N-litter whereas those that are positioned towards the right (blue) incorporated relatively more isotope from ^13^C-exudates. Probes targeting protists and nematodes were on the chip (Table S1) but we did not measure any significant enrichment in this 3-day ^13^CO_2_ experiment.

## DISCUSSION

While it is widely recognized that fungi and soil fauna are instrumental to decomposition (6), less is known about how decomposition processes interact with the greater soil food web, which includes bacteria, archaea, fungi, and microfauna. To this end, we directly sequenced total RNA to identify bacteria, archaea, and eukaryotes in the presence and absence of root litter, and determined how the presence of a living root altered these communities. We also used NanoSIMS-enabled microarray analysis (Chip-SIP) to track the fate of ^15^N root litter and ^13^C rhizodeposits and identified the substrate preferences of abundant organisms in the rhizosphere amended with litter.

### Protists were abundant in decomposing litter and are important for nutrient cycling

We found that protists were abundant in decomposing litter in the presence and absence of living roots. In soil, decomposition is carried out by bacteria, fungi, mesofauna (e.g. microinvertebrates), and macrofauna (e.g. earthworms, millipedes), whose shredding action create smaller particles that are more readily accessible to microbes (6). On the other hand, soil microfauna (e.g. protists, nematodes) primarily consume bacteria and fungi (67–69). Those that feed directly on soil bacteria and fungi are thought to indirectly influence the decomposition of soil organic matter in multiple ways. For example, not only can microfauna alter the composition of bacterial and fungal decomposers, they can also accelerate the turnover of microbial biomass and excrete nutrients derived from microphagy (12, 70), which in turn can enhance litter decomposition (24). Protists can stimulate microbial nutrient cycling through the microbial loop (71, 72), which is a phenomenon where N contained in microbial biomass is higher than the N demand of protists, and predation ultimately leads to an increase in available N after excretion.

The high abundance of protists we observed in decomposing litter may have altered the flow of nutrients though grazing and is an important dynamic currently missed in many bacterial and fungal litter decomposition studies.

### Amoebas are potentially important yet overlooked top-down driver of detritusphere community dynamics and nutrient cycling

Rhizaria (Cercozoa), Amoebozoa, and Alveolata were the most abundant protists in our soils. A previous metatranscriptomic study also found that Rhizaria and Amoebozoa were abundant in grassland soils relative to peatlands (39). Interestingly, we predominantly observed Rhizaria in unamended bulk soil, whereas Amoebozoa were more abundant when litter was present in both bulk soil and the rhizosphere. This suggests these two abundant groups inhabit different niches within the soil environment. In a complementary transcriptomics dataset from this soil (52), Amoebozoa expression of exoproteases was highest in the litter-containing rhizosphere and bulk soils (Figure S2), which further suggest these organisms play an active role in microbial community dynamics in the detritusphere. Microfaunal predation is generally overlooked as a top-down driver of microbial community assembly (73). Amoebas in particular are known to be mycophagous or bacterivores (15, 16) and can influence microbial community structure (12). As the Amoebozoa Supergroup is typically missed in amplicon analysis (74), our results suggest Amoebas may be an overlooked contributor to microbial community dynamics and nutrient cycling in the detritusphere.

### Rhizosphere and bulk soil harbor unique litter-decomposing communities

Each treatment in this study had a unique microbial community, where the rhizosphere and bulk soil contained distinct assemblages of potential decomposers. In particular, the bulk-litter community had significantly more Bacteroidetes, Proteobacteria (Oxalobacteriaceae, Caulobacteraceae, Bacteriovoraceae), and saprotrophic *Chaetomium* fungi relative to the unamended control, but was depleted in Actinobacteria and Acidobacteria. In the rhizosphere, both sequencing and ^15^N-litter Chip-SIP identified members of the Sphingobacteriaceae as responders to litter; Chip-SIP also identified Xanthobacteriaceae bacteria and Eurotiomycetes fungi as consumers of ^15^N-litter in the rhizosphere. Previous stable isotope probing studies have identified taxa from the Actinobacteria, Firmicutes, and Bacteroidetes as bacterial decomposers of plant material in soil (75, 76), and taxa from the Burkholderiales, Caulobacteriales, Rhizobiales, Sphingobacteriales and Xanthomonadales as cellulose-degrading bacteria (77). The low abundance of Actinobacteria in the bulk-litter soil was unexpected given that they are commonly identified as soil decomposers (75–77), and the trend toward reduced total fungi transcripts (Fig. 4b) did not support our initial hypothesis. In bulk-litter soil, a different suite of decomposers may have preferred the root litter in our system (e.g., Bacteroidetes, *Chaetomium* fungi) instead of Actinobacteria. Alternatively, the reduction in Actinobacteria and trend toward reduced total fungi in the presence of litter might be explained by interactions among soil microbes. As Amoebozoa and bacterial predators (e.g., Bacteriovoraceae) were abundant in the litter-containing treatments, it is possible that the low abundance of common decomposers in the presence of litter might have been driven by fungal or bacterial grazing (78, 79). Members of the Oxalobacteriaceae have also been shown to consume fungi (80), though we did not identify any known mycophageous genera in our dataset. We also note that we did not observe any macrofauna or mesofauna sequences in the dataset; macrofauna (and possibly mesofauna) would have been removed during the processing phase of soils, or because the amount of soil extracted for DNA was too small in volume.

### Isotopic tracers identify substrate preferences in the rhizosphere

The Chip-SIP results helped us to start to disentangle microbial substrate preferences in the rhizosphere. We used two isotopic tracers to determine if soil microbes preferentially consumed ^13^C-exudates or ^15^N-litter. In the rhizosphere-litter treatment, the Actinobacteria tended to prefer ^15^N-litter, while Fungi and the Rhizobiales tended to assimilate ^15^N-litter and ^13^C-exudates equally, with a slight preference for ^13^C-exudates. While Actinobacteria were the second-most abundant phylum in the rhizosphere treatments, the Chip-SIP data indicated that they may have preferred to consume detrital SOM in this habitat; this is consistent with recent findings where the Actinobacteria had the highest CAZyme gene expression in the detritusphere or aging rhizosphere (rhizosphere >20 days old) (52). Actinobacteria have been identified as plant-or cellulose-degraders by density gradient SIP (75–77). This highlights that 16S patterns alone have a limited ability to determine substrate preferences and require additional information to assess microbial ecophysiology, such as through isotope tracing or activity-based analyses.

### Relevance of the microbial food web for soil C cycling

The multi-trophic changes we observed in our study are relevant for soil carbon cycling. Microbial communities have diverse arrays of physiological strategies for consuming carbon substrates (7). Changes in the microbial community can influence the rate of decomposition and decomposition products (7, 81), and are expected to alter the diversity of compounds available for sorption to mineral surfaces (82). For example, we found that Actinobacteria decreased in the bulk-litter treatment while *Chaetomium* fungi increased, which likely altered the composition of exoenzymes available to breakdown plant material and can further influence the rates and types of organic matter that can be degraded (83, 84). Microbial bodies themselves define the forms of C available for stabilization, such as cell wall components, polysaccharides, enzymes, intracellular sugars, and intermediate products of litter decomposition (7, 83, 85-87); these components can have different residence times in soil (83, 88). Selective predation, such as by protists or Bacteriovoraceae, can alter the both the taxonomic and functional composition of the soil microbiome (73). In addition, predation by protists potentially diversifies and alters forms of C available for sorption to mineral surfaces since the microbial polymers undergo digestion before excretion, and protists have been shown to selectively retain particular classes of metabolites (89). Thus, the trajectory of carbon through the soil food web likely impacts the fate and persistence of carbon in soil.

## CONCLUSIONS

Using metatranscriptomic sequencing of total RNA, we identified the bacterial, archaeal, and eukaryotic communities that responded to detrital root litter in rhizosphere and bulk soil. Litter-decomposing communities differed depending on the presence and absence of a root, where the litter-amended bulk soil had the most distinct microbial and protist communities. The litter-amended rhizosphere and bulk soil contained significantly more Amoebozoa than the unamended controls, and grazing by these protists may be an important top-down control on the microbial community during litter decomposition and alter the trajectory of carbon through the soil food web. Chip-SIP NanoSIMS analysis identified substrate preferences for ^15^N root detritus or ^13^C rhizodeposits fresh root exudates and gave insights into food web processes that were not discernible from compositional analyses alone. Future work combining shotgun RNA community analyses and stable isotopes has the potential to improve our ability to track nutrients through multi-trophic food webs.

## ACKNOWLEDGEMENTS

This research was supported by the U.S. Department of Energy Office of Science, Office of Biological and Environmental Research Genomic Science program under awards SCW1421, SCW1589, and SCW1678 to JPR and under awards DE-SC0010570, DOE-SC0016247, and DE-SC0020163 to MKF. Work conducted at Lawrence Livermore National Laboratory was supported under the auspices of the U.S. DOE under Contract DE-AC52-07NA27344. Work conducted at Lawrence Berkeley National Laboratory was supported under Contract DE-AC02-05CH11231. We thank Katerina Estera Molina and Shengjing Shi for laboratory assistance, Donald Herman for assistance with the EPIC ^13^CO_2_ labeling system, Ulas Karaoz for data management assistance, and Ella Sieradzki for the Acanthamoebidae supplemental figure.

## SUPPLEMENTAL

### Production of ^15^N-labeled root litter

To create ^15^N-labeled root litter, *Avena fatua* was grown for 8 weeks in fritted clay and fertilized solely with a custom mixture of Hoagland’s plant nutrient solution, where the nitrogen-containing compounds were replaced with their 99 atom % ^15^N analogs (0.505 g/L K^15^NO_3_ (Cambridge Isotope Laboratories), 0.59 g/L Ca(^15^NO_3_)2 x 4H_2_O (Cambridge Isotope Laboratories), 0.0225 g/L Sprint 330 iron chelate (Becker Underwood), 0.493 g/L MgSO_4_ x 7H_2_O, 0.080 g/L ^15^NH_4_^15^NO_3_ (Cambridge Isotope Laboratories), 2.86 g/L H_3_BO_3_, 1.81 g/L MnCl_2_ x 4H_2_O, 0.22 g/L ZnSO_4_ x 7H_2_O, 0.051 g/L CuSO_4_, 0.09 g/L H_3_MoO_4_ x H_2_O, and 0.5 ml/L of 1M KH_2_PO_4_ (pH 6.0)). Roots were triple washed in deionized water, dried, and stored for 1 year prior to use. Roots were chopped to ca. 1 mm lengths using scissors.

### Data processing using EMIRGE

Since EMIRGE probabilistically reconstructs sequences, the near full-length sequences have a higher error rate than sequences directly generated by Illumina sequencing. To address this, singletons and sequences present in only one sample were removed from the dataset. Next, we assessed the efficacy of three chimera checking tools to identify potential chimeras created by EMIRGE (UCHIME (62), DECIPHER (90), Chimera Slayer (91)), and determined that UCHIME was the most effective chimera-checking tool for our dataset. To assess the chimera-checking tools, we used a set of EMIRGE sequences reconstructed from a mock community composed of 52 known isolates available as a supplemental dataset from Miller, Baker (36). Any novel sequences in this EMIRGE dataset are likely chimeras, since the SSU sequences of all the community members are known. We determined that 3 of 23 reconstructed sequences were putative chimeras, since they were < 90% similar to organisms in the NCBI database using BLAST. We tested each chimera-detection method to determine if it could identify these 3 sequences in the EMIRGE dataset. Only UCHIME was capable of detecting chimeras in the dataset; it was able to identify 2 of the 3 putative chimeras. Therefore, UCHIME was used for chimera-checking analyses.

### Probe design

We designed a custom Chip-SIP array for rhizosphere soil with probes targeting bacteria, archaea, and eukaryotes (fungi, protists, nematodes). Probes were designed from the sequences reconstructed by EMIRGE from the four treatments in this experiment. The 180 most abundant OTU from bacteria, archaea, and eukarya were targeted for probe design. The microarray probes were designed in ARB using the following restrictions: (≤ 2 mismatches tolerated, GC content < 80%, homopolymer runs ≤ 4bp) (65). Twenty-five different probes were selected for each sequence that were unique relative to the SILVA database and RNA-Seq databases. Based on preliminary fluorescence data (using soils from actual experiment), from these 25 probes, we selected the 10 probes that had the highest hybridization scores to synthesize on the final microarray (signal:noise ≥ 1.3). Sequences that had few probes with positive fluorescence (signal:noise < 1.3) were added to the microarray by keeping 10 probes with the best ARB score (a measure of how specific the probe is to the sequence of interest).

### Chip-SIP microarray synthesis

Microarrays were coated with a conductive surface prior to probe synthesis to eliminate charging during SIMS analysis. Glass slides coated with indium-tin oxide (Sigma) were treated with an alkyl phosphonate hydroxy-linker to provide a starting substrate for probe synthesis (92). Microarray probes (spot size = 17 μm) were synthesized using a photolabile deprotection strategy (93) on a Nimblegen Maskless Array Synthesizer (Roche). The probe sets were laid out in horizontal lines across the chip. All the probes were printed three times in three replicate blocks on the microarray. Nimblegen synthesis reagents (Roche) were delivered through the Expedite system (PerSeptive Biosystems).

### Microarray hybridization

Two microarrays are necessary for Chip-SIP: a standard fluorescence microarray and a separate NanoSIMS microarray. Fluorescence analysis is necessary to confirm that the probes are hybridized with RNA. However, labeling RNA with a fluorophore introduces ^12^C-carbon that dilutes the ^13^C signal. Therefore, the RNA samples were split for fluorescence and NanoSIMS analyses and the RNA used in the NanoSIMS analysis was left unlabeled. Microarrays were not replicated.

For fluorescence analysis, the RNA was labeled with Alexafluor 532 dye using the Ulysis kit (Invitrogen), and were incubated for 10 min at 90°C (2 μL RNA, 10 μL labeling buffer, 2 μL Alexafluor reagent) and subsequently fragmented. RNA for NanoSIMS analysis was not labeled, and instead preceded directly to fragmentation. Samples were fragmented using 5X fragmentation buffer (Affymetrix) for 10 min at 90°C. Fragmented RNA was purified using a Spin-OUT™ minicolumn (Millipore), and RNA was concentrated by ethanol precipitation to a final concentration of 500 ng μL^-1^. For array hybridization, RNA samples were mixed with 1X Hybridization buffer (Nimblegen) and placed in a Nimblegen X4 mixer slide. The arrays were incubated inside a Maui hybridization system (BioMicro Systems) for 18 hrs at 42°C and then washed according to manufacturer’s instructions (Nimblegen).

Arrays with fluorescently labeled RNA were imaged with a Genepix 4000B fluorescence scanner at pmt = 650 units. Arrays with non-fluorescently labeled RNA were marked with a diamond pen and also imaged with the fluorescence scanner to subsequently navigate to the analysis spots in the NanoSIMS. Slides were trimmed and mounted in custom-built stainless-steel holders.

### Chip-SIP NanoSIMS analyses

Mass resolution was set to ~10,000 mass resolving power to minimize the contribution of isobaric interferences to the species of interest (e.g., so that ^11^B^16^O^-^ contributes < 1/100 of the ^13^C^14^N^-^ ratio, and ^13^C_2_^-^ contributes < 1/1000 of the ^12^C^14^N^-^ ratio). Analyses were performed in imaging mode to generate digital ion images of the microarray for each ion species. The primary beam current was 5 to 7 pA Cs^+^, which yielded a spatial resolution of 200-400 nm and a maximum count rate on the detectors of ~300,000 cps ^12^C^14^N. Analysis area was 50 x 50 μm^2^ with a pixel density of 256 x 256 with 0.5 or 1 ms/pixel dwell time. Ion counts were corrected for detector dead time on a pixel-by-pixel basis.

### Chip-SIP statistical analyses

Individual 50 x 50 μm^2^ isotope ratio images were stitched together to create an isotope map of the microarray surface using custom software developed for NanoSIMS analysis (L’image, L. Nittler, Carnegie Institution of Washington). Probe spot regions of interest (ROIs) were selected by hand or with an autodefinition function, and ^15^N/^14^N and ^13^C/^12^C isotope ratios were calculated for each ROI. Isotope ratios were converted to atom percent excess (APE) values using the formula APE = [R_meas_/(1 + R_meas_) – R_control_/(1 + R_control_)] × 100%, where R_meas_ is the isotope ratio measured by NanoSIMS and Rcontrol is the mean ^15^N APE or ^13^C APE value for the control probe locations. The presence of RNA on each probe was confirmed using a separate fluorescence microarray analysis, where hybridized probes had a signal to noise ratio > 1.3.

Two criteria identified which OTUs were enriched with ^13^C or ^15^N. First, each probe set was required to have five of ten probes with a signal:noise ratio > 1.3; this determined which OTUs were present in the dataset. Second, each probe set had to have five probes with enrichment of either >0.020 ^13^C atom percent excess (APE) or >0.011 ^15^N APE (equivalent to 30‰ for both). For the OTUs that met these criteria, we calculated a ^13^C-APE:^15^N-APE ratio for the entire probe set by averaging all APE enrichments > 0 for ^13^C and ^15^N and then dividing the two averages (average ^13^C APE / average ^15^N APE).

**Figure S1.**
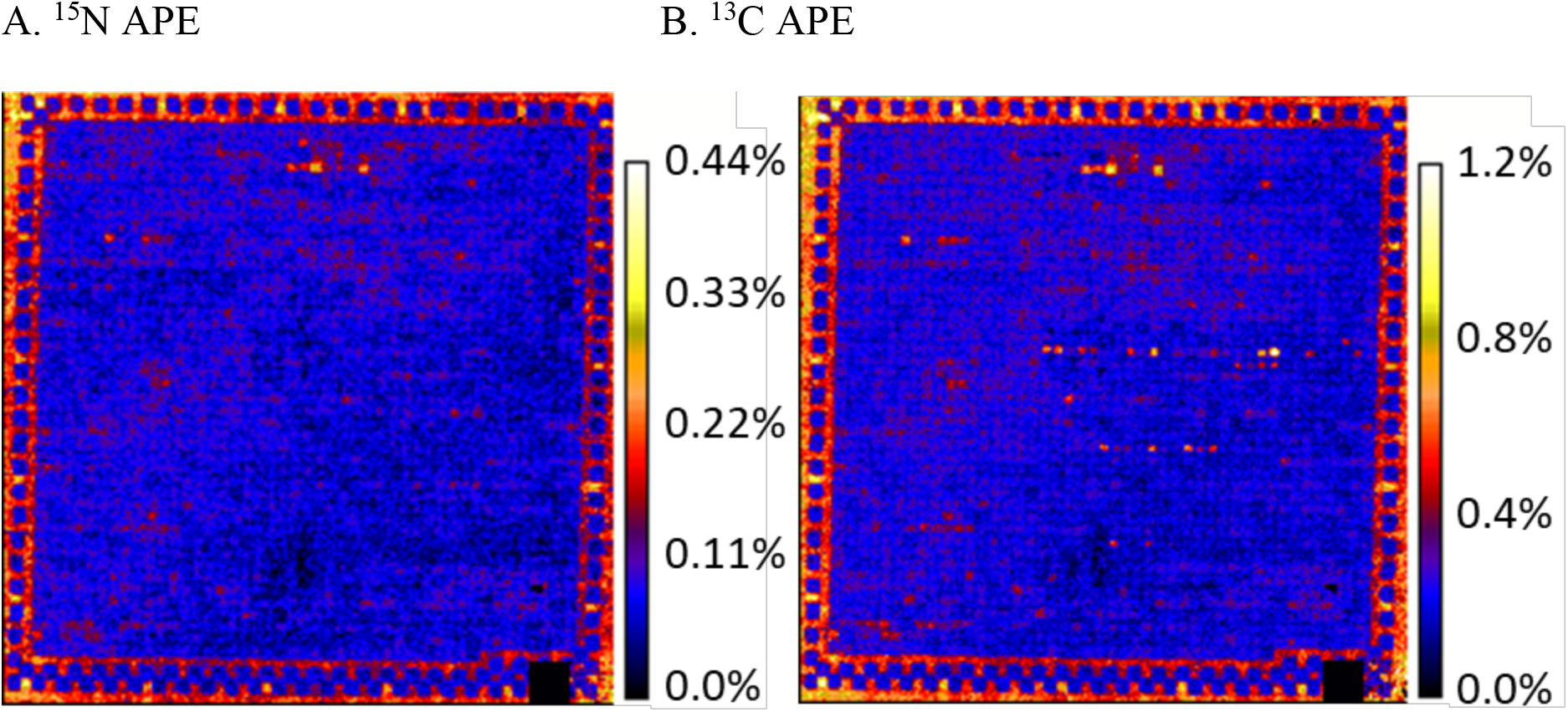
Chip-SIP isotope maps of a single phylogenetic microarray hybridized with isotopically labeled RNA from a rhizosphere soil microbial community exposed to ^15^N-labeled root detritus and ^13^C-exudates. As the microbes consume the substrates with the different isotope labels, they assimilate the isotopes into their microbial biomass and nucleic acids, and their preference for the ^15^N-root litter or ^13^C-exudates is determined by the amount of (A) ^15^N and (B) ^13^C contained in the RNA hybridized to a probe set specific to each taxon. Color scale bars indicate atom percent excess (APE) enrichment of the microarray surface. Probes sets are arranged in horizontal lines on the chip. The brightest ^13^C-enriched probe sets with no visible corresponding ^15^N-enriched probes are from plant host RNA.

**Figure S2.**
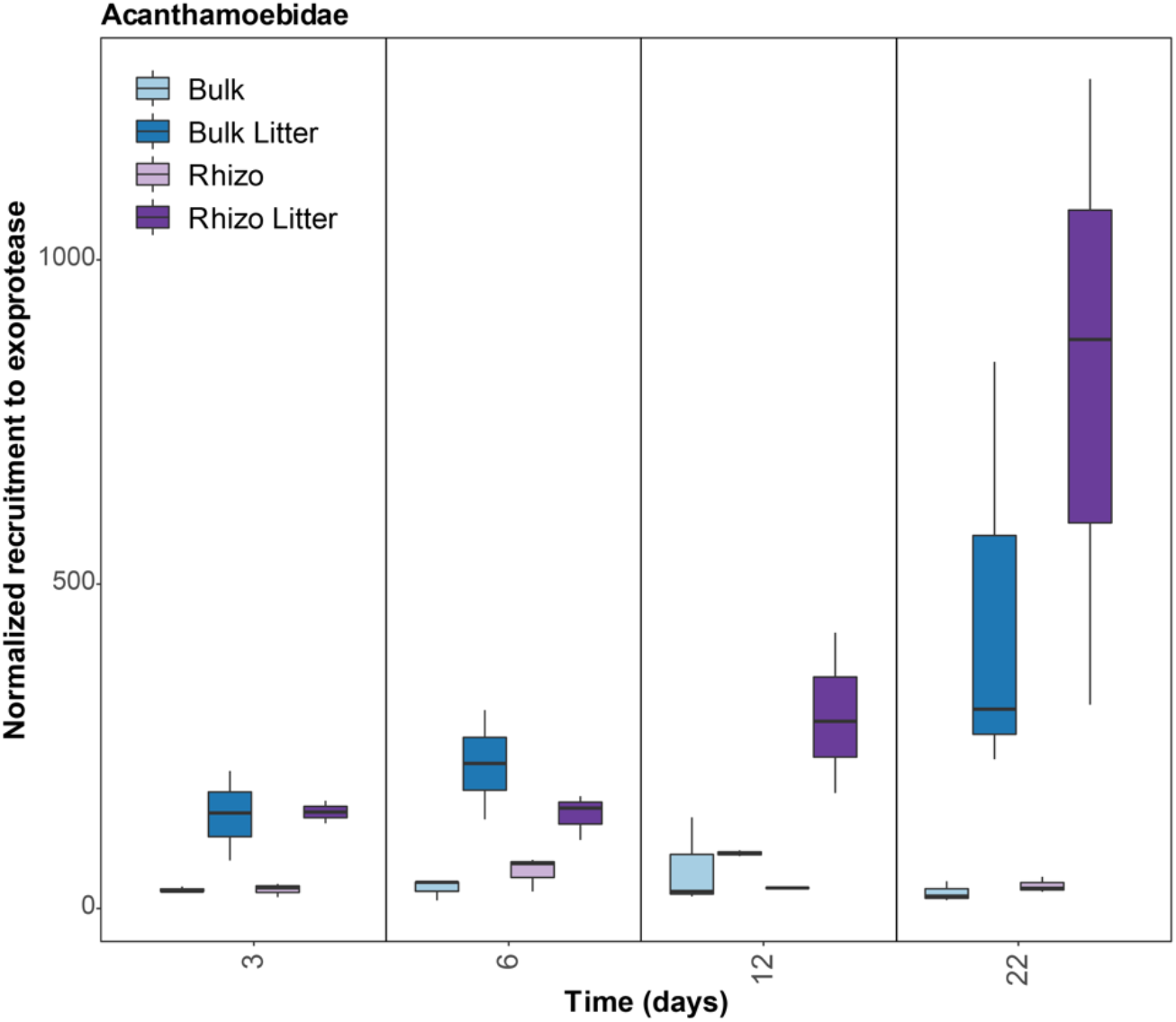
Acanthamoebidae exoprotease gene expression from a complimentary transcriptomic dataset on the same soil (52) from 3-22 days. Rhizosphere and bulk soil were amended with detrital root litter (Rhizo Litter, Bulk Litter) or unamended (Rhizo, Bulk). Sequences were normalized using DESeq2 as previously (52).

